## Supplementary figures for "A primate-specific retroviral enhancer wires the *XACT* lncRNA into the core pluripotency network in human"

**Supplementary figure 1 – The transcriptional landscape of the *XACT*/*T113.3* locus is different across primate species**

**A.** International Human Epigenome Consortium (IHEC) strand-specific RNA-seq from human, chimpanzee and rhesus macaque iPSCs across the region spanning the *LHFPL1* and *LRCH2* genes[1,2]. Schematic representation of the genes found across the locus are shown below each panel. Syntenic regions for the human *XACT* and *T113*.3 genes are shown for rhesus macaque as grey rectangles.

Supplementary figure 1

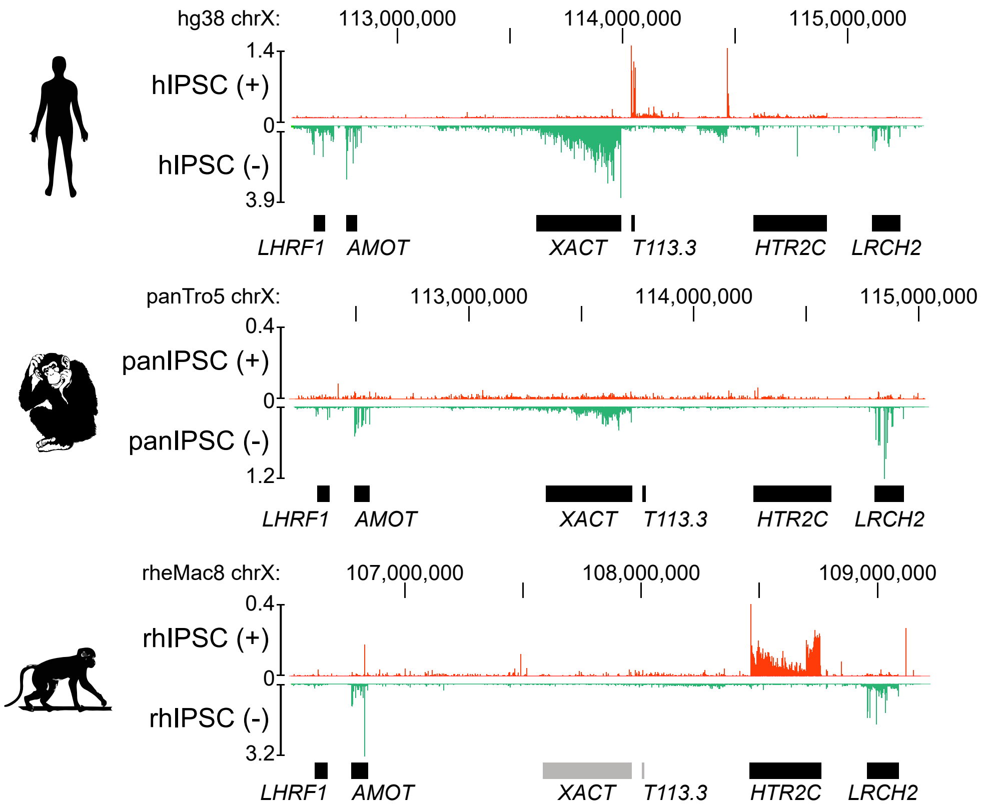

**Supplementary figure 2 – *XACT*, but not *XIST*, shows a similar expression dynamics to *T113.3* in pluripotent contexts**

**A.** Analysis of *XACT* (green) and *T113.3* (red) expression by RNA-FISH in female H9 hESCs. Percentage of cells co-expressing *XACT* and *T113.3* from the same locus is indicated. Quantification of the different patterns of expression is represented on the right (n=193). The white scale bar represents 5 µm.

**B.** Quantitative RT-PCR analysis of *XACT* and *T113.3* expression during a 10-day undirected differentiation of female WIBR2 hESCs. The bar charts correspond to the average of two independent differentiation experiments. Error bars indicate the s.d. The correlation coefficient for the expression dynamics of the two genes is shown (r= 0.97, Pearson correlation).

**C.** Boxplot of *XACT* and *T113.3* expression levels (log2 RPKM) according to developmental stage. Dataset from[3–5].

**D.** Plot of *T113.3* versus *XIST* expression levels (log2 reads per kilobase per million mapped reads [RPKM]) in early (E3, E4 and early E5) and late stage (E5, E6 and E7) female (upper panels) and male cells (lower panels), with corresponding Spearman correlation score and p-value. Dataset from[6].

Supplementary figure 2

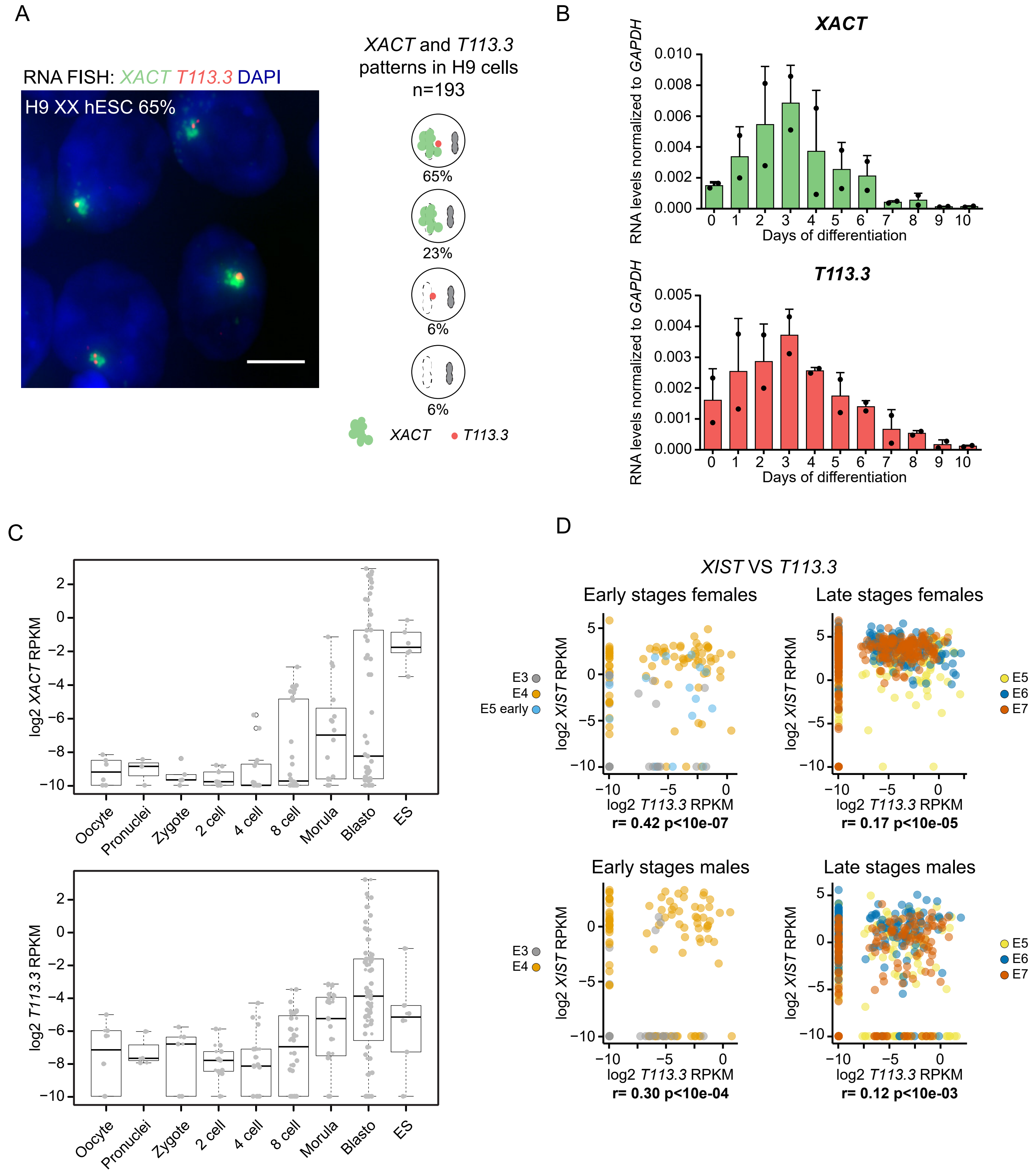

**Supplementary figure 3 – Interfering with the transcription of *XACT* or *T113.3* does not impact on expression of pluripotency markers**

**A.** Quantification of *NANOG* and *OCT4* steady state RNA levels by quantitative RT-PCR upon *XACT* CRISPRi (left bar charts) and *T113.3* CRISPRi (right bar charts) in H1 hESCs.

**B.** ChIP-qPCR analysis of H3K4me3 (top graphs) and H3K27ac (bottom graphs) enrichment across the *XACT*/*T113.3* locus upon *XACT* CRISPRi (green squares) or *T113.3* CRIPRi (red circles). The positions analysed are indicated with arrows. ChIP-qPCR data for untransfected cells (gray inverted triangles) and cells transfected with a vector without a sgRNA (black triangles) are also shown. Right graph represents the relative enrichment of each mark in control positions (SOX2 TSS, B2M TSS and hXIC19). Statistical significances were determined using a 1-way ANOVA test (all conditions compared to sg_empty). P-values: < 0,05 (*), <0,01 (**), <0,001 (***) and <0,0001 (****).

Error bars indicate the s.d (minimum of 3 replicates per condition).

Supplementary figure 3

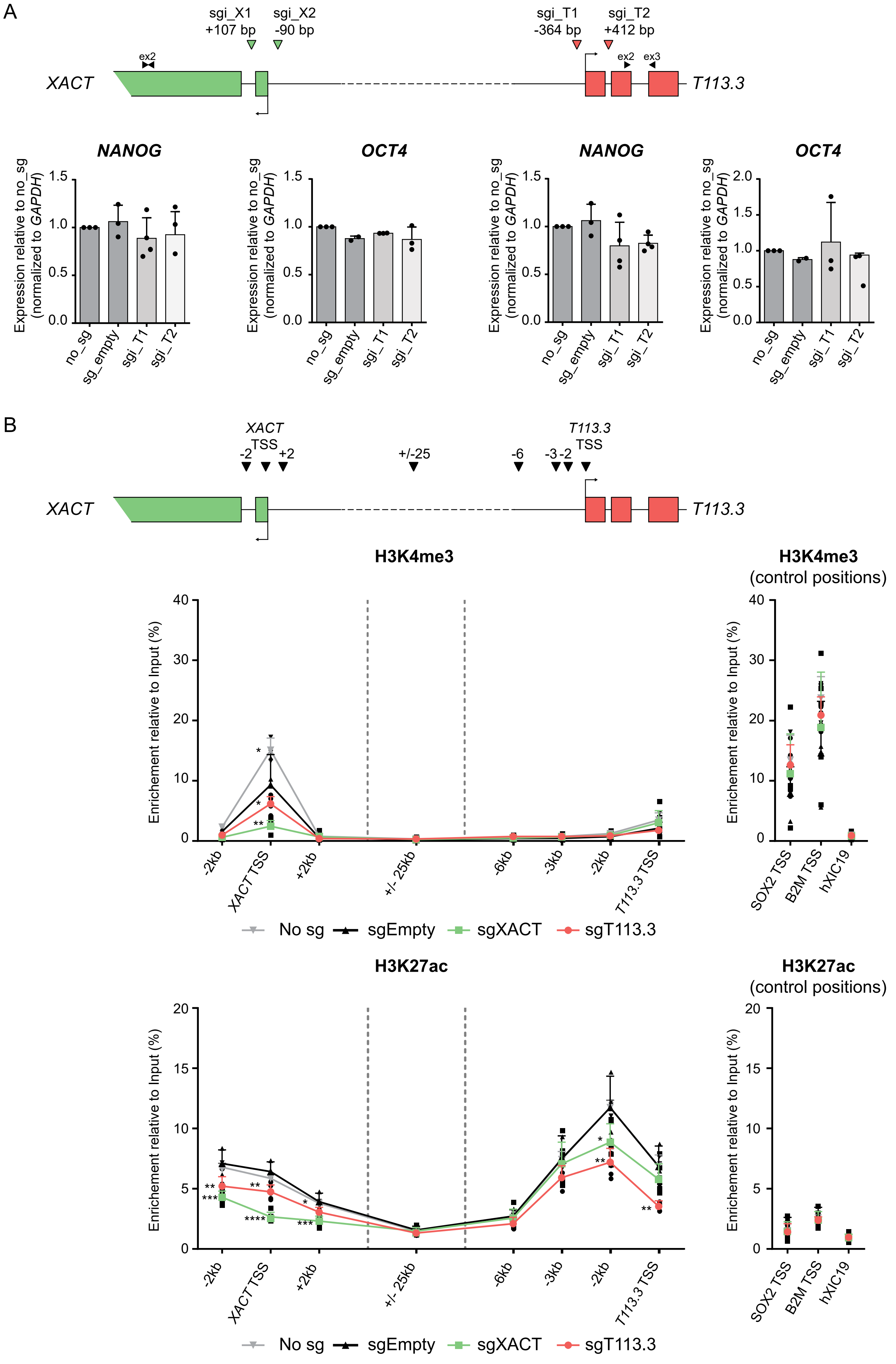

**Supplementary figure 4 – The LTR7/HERVH *T113.3* is dispensable for the transcriptional regulation of the *XACT*/*T113.3* locus in pluripotent cells and during undirected differentiation of H1 hESCs**

**A**. Quantification of the global RNA levels of *OCT4* (left bar chart) and *NODAL* (right bar chart) by RT-qPCR upon *XACT* LNA gapmers or *T113.3* LNA gapmers KDs. The bar charts correspond to the average of at least 5 replicates per condition.

**B**. Schematic representation of the *T113.3* deletion strategy (left panel), showing the positions of the CRISPR sgRNAs (red triangles) and screening primers used to characterize the clones (black arrows). The right panels represent electrophoretic gel pictures showing the PCR amplicons obtained with the different screening primers on genomic DNA from the different clones. Top panel: PCR assay detecting the deleted allele (PCR 1-2); middle panel: PCR assay detecting the inversion of the *T113.3* gene (PCR 3-4); bottom panel: PCR assay testing the presence of the *T113.3* gene (found in WT or INV clones). Selected clones are highlighted in red.

**C**. Quantification of the steady state RNA levels of *OCT4* (pluripotency marker), *NODAL* (differentiation marker), *AMOT*, *HTR2C* (genes flanking the *XACT*/*T113.3* locus) and *ATRX* (gene on the X chromosome) by quantitative RT-PCR in H1 *T113.3* WT, KO and INV clones. The bar charts correspond to the average of two or three independent clones.

**D.** Quantitative RT-PCR analysis of *T113.3*, *OCT4*, *HAND1* and *GATA6* expression during a 10-day undirected differentiation of H1 T113.3 WT, KO and INV hESC lines. $n \geq 2$.

Error bars indicate the s.d.

Supplementary figure 4

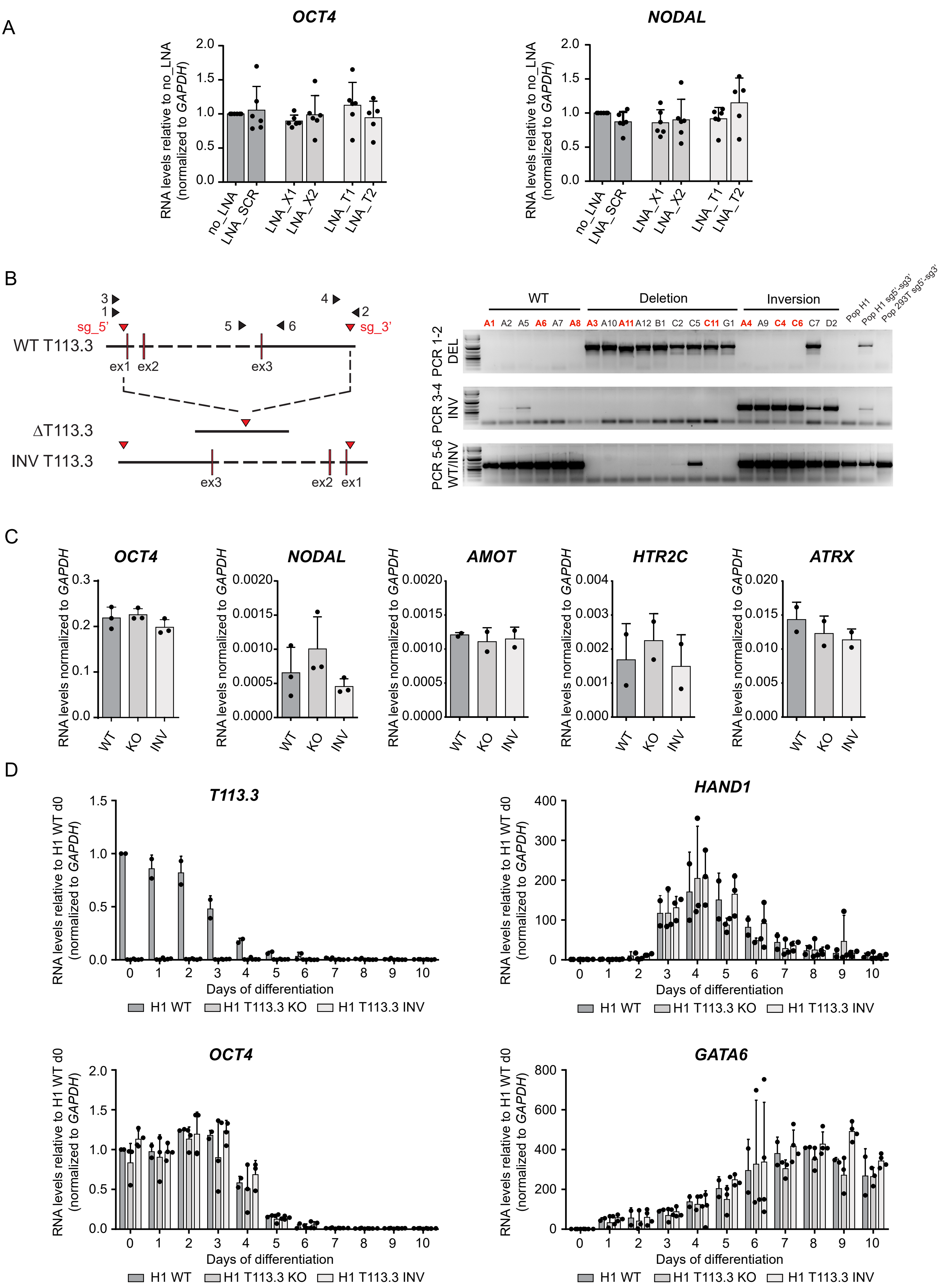

**Supplementary figure 5 – The LTR/ERV element upstream of *T113.3* is a pluripotent specific enhancer**

**A**. ATAC-seq (black) and ChIP-seq data for H3K4me3, H3K27ac (blue), CTCF (brown), OCT4, SOX2 and NANOG (orange) enrichment across the *XACT*/*T113.3* locus in H1 hESCs (top panel). ChIP-seq data for H3K4me3 and H3K27ac enrichment in IMR90 is shown below. Publicly available data was obtained from the ENCODE project[7] and CISTROME DB[8].

**B.** Schematic representation of the *T113.3* locus, showing the positions of the CRISPR sgRNAs (red triangles) used for targeting of the CTCF (top scheme) or TFB sites (bottom scheme) and the primers used for genotyping (black arrows). Panels bellow each scheme represent electrophoretic gels showing the PCR amplicons obtained with the different screening primers on genomic DNA from H1 CTCF- (top) and TFB- (bottom) targeted clones. Top panels: PCR assay detecting the deleted and WT alleles (PCR 1-3 for CTCF site and PCR 4-6 for TFB site); bottom panel: PCR assay detecting inversion of the targeted region (PCR 1-2 for CTCF site and PCR 5-6 for TFB site).

**C.** Quantification of *OCT4* (left bar chart) and *NANOG* (right bar chart) steady state RNA levels by RT-qPCR in CTCF- (top bar charts) and TFB- (bottom bar charts) targeted H1 clones. The bar charts correspond to the average of at least, two independent clones.

**D**. RT-qPCR analysis of the RNA levels of *XACT* (top left bar chart), *T113.3* (top right bar chart), *GATA6* (bottom left bar chart) and *PAX6* (bottom right bar chart) in cells transfected with siRNA targetting OCT4, SOX2, NANOG or different combinations of these siRNAs. Bar charts correspond to the average of five independent experiments. Statistical significances were determined using a 1-way ANOVA test (all tests compared to siSCR). P-values: < 0,05 (*), <0,01 (**), <0,001 (***) and <0,0001 (****).

Error bars indicate the s.d.

Supplementary figure 5

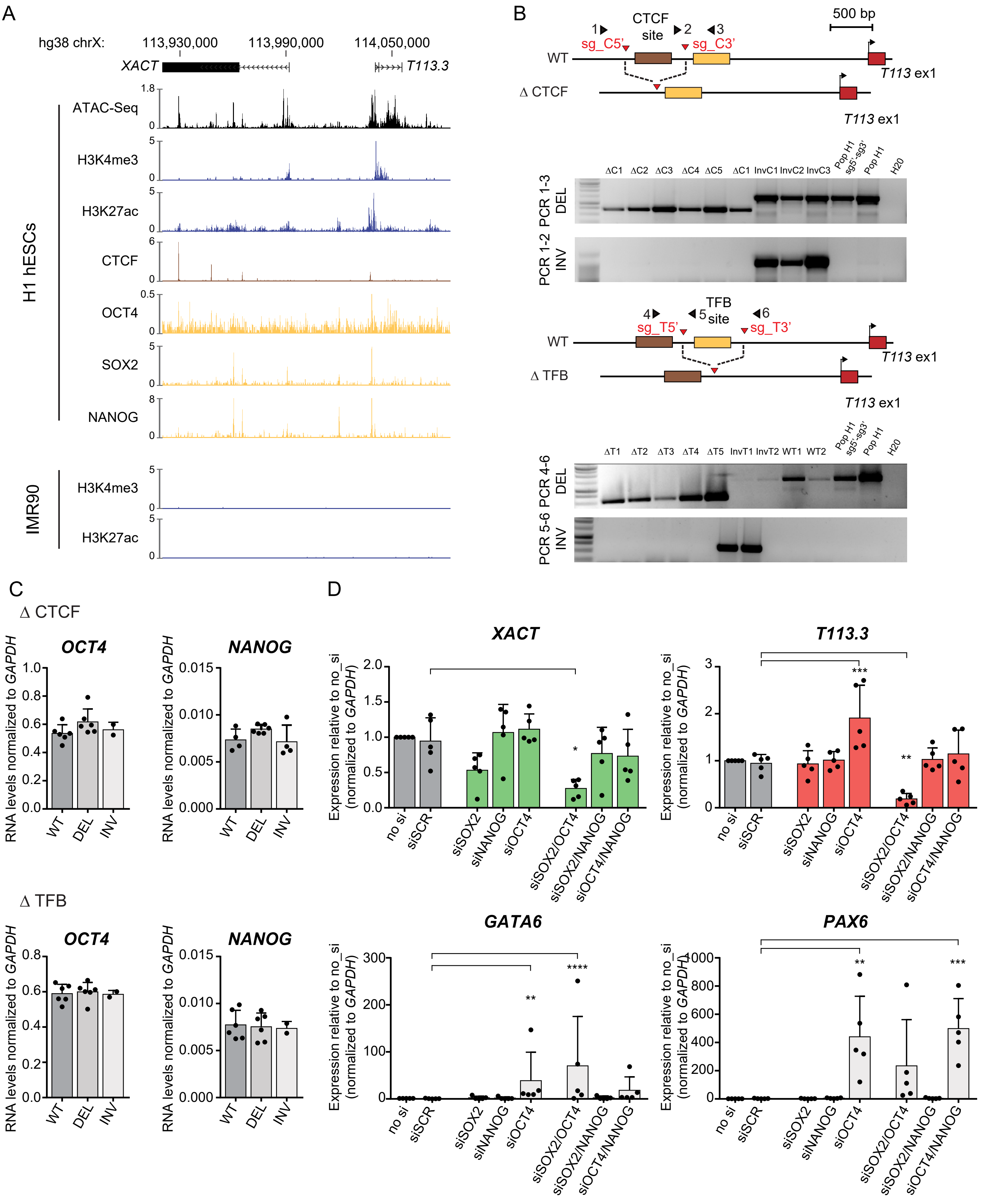

**Supplementary figure 6 – The LTR48B enhancer is active in species that do not have *XACT* and *T113.3* and evolved independently of the core of the LTR48B subfamily**

**A.** ChIP-seq for H3K27ac and strand-specific RNA-seq from human (left panel) and rhesus macaque (right panel) iPSCs over a region spanning the *XACT* promoter and *T113.3* gene. Schematic representation of the *XACT* and *T113.3* genes as well as their common enhancer is shown below each panel. Dataset from[1].

**B.** Comparison between the *XACT*/*T113.3* LTR48B enhancer and all the other human LTR48B elements. Multiple alignment was performed between the LTR48B enhancer and the consensus sequence for the LTR48B family. P-values for the likelihood of the consensus sequence being a binding site for pluripotency transcription factors were determined using RSAT matrix-scan. The logo of all the human LTR48B sequences was obtained from the Dfam website[9].

**C.** Heatmap of log2-normalized ChIP-seq signal for OCT4 and NANOG in hESCs. The heatmaps show the distribution of the transcription factors in 8 kb windows (4 kb in each direction) centred at the middle of LTR7 (left panels) and LTR48B elements (right panels). The summary plot representing the average coverage of the transcription factors across the 8 kb windows around LTR7 (left) and LTR48B (right) are represented above the heatmaps. Dataset was from[10].

**D.** LTR7/HERVHs elements are enriched in OCT4 and NANOG pluripotency factors. Boxplot represents normalized number of reads (log2 reads per kilobase per million mapped reads [RPKM]) that mapped to individual LTR elements belonging to either LTR7 or LTR48B subfamilies. A thousand random regions across the genome were selected to use as a randomized control. p-values were calculated from a t-test. Dataset was from[10].

**E.** Number of LTR7 (left diagram) and LTR48B (right diagram) elements that are bound by OCT4 (red) and/or NANOG (blue) in hESCs.

Supplementary figure 6

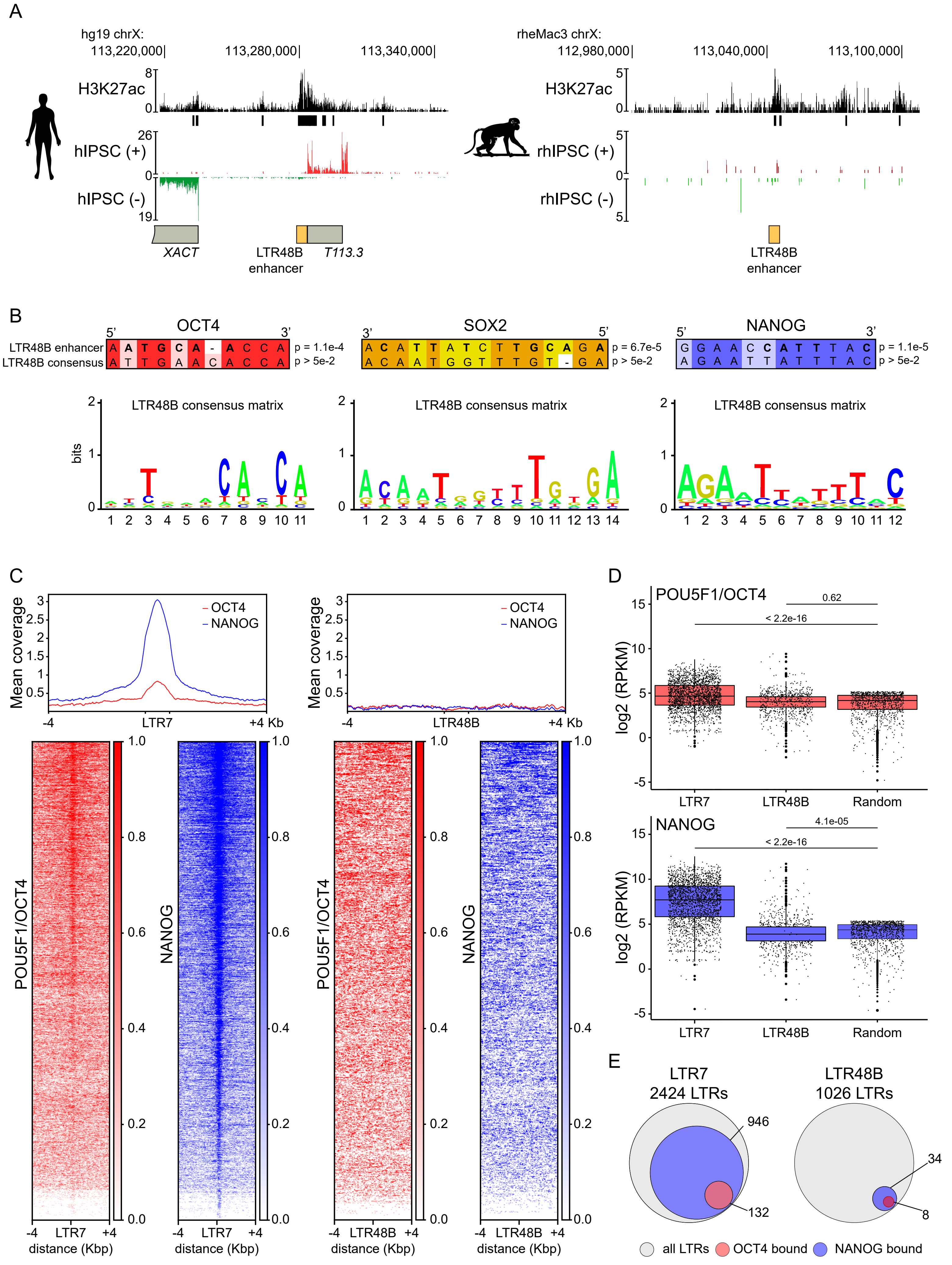
